## Supplementary figures and images for "Comparison of SVM and Spectral Embedding in Promoter Biobricks’ Categorizing and Clustering"

### Supplemental Fig 1

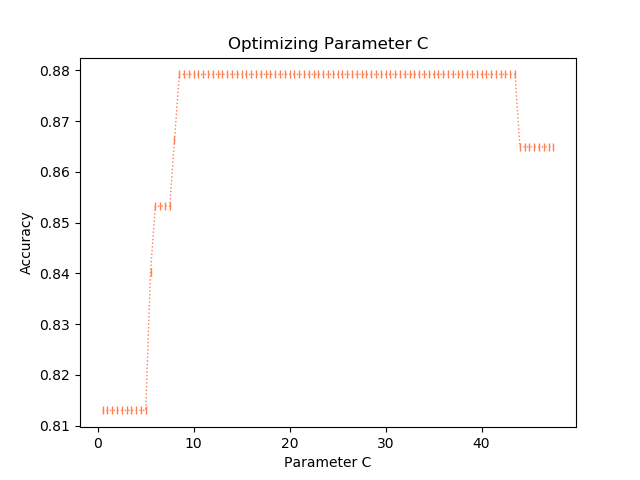

### Supplemental Fig 2

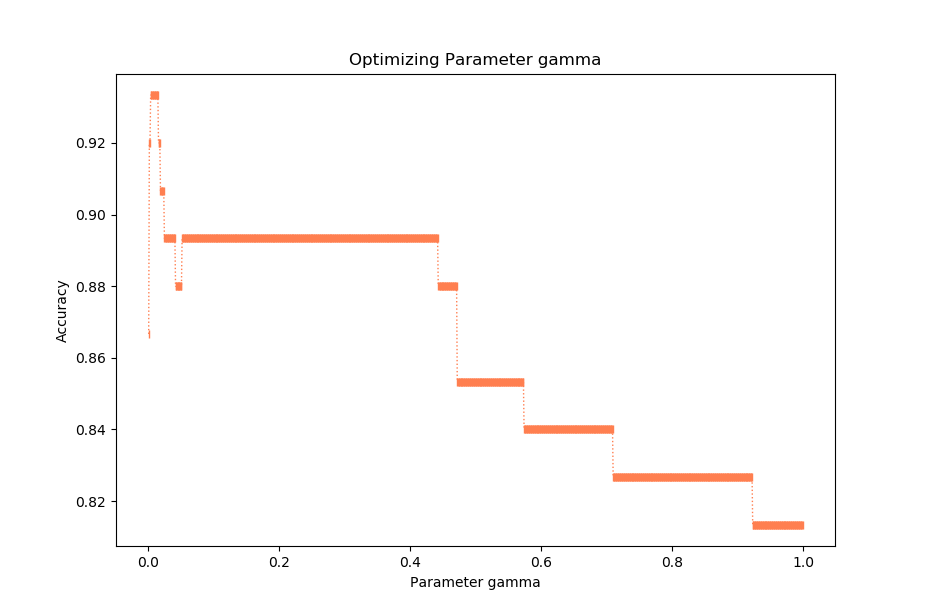
